# Widespread symbiosis of ciliate epibionts colonizing gills of shrimps inhabiting vents and seeps across the Pacific Ocean

**DOI:** 10.64898/2026.02.27.708428

**Authors:** Lison Hey, Chong Chen, Ting Xu, Emily Cowell, Dewi Langlet, Pierre Methou

**Affiliations:** Univ Brest, Ifremer, BEEP, F-29280 Plouzané, France; X-STAR, Japan Agency for Marine-Earth Science and Technology (JAMSTEC), 2-15 Natsushima-cho, Yokosuka, Kanagawa 237-0061, Japan; Southern Marine Science and Engineering Guangdong Laboratory (Guangzhou), Guangzhou, China; Department of Ocean Science and Otto Poon Center for Climate Resilience and Sustainability, The Hong Kong University of Science and Technology, Hong Kong, China; Department of Biology, Temple University, Philadelphia, PA, USA; Institut Systématique Evolution Biodiversité (ISYEB), Muséum National D’Histoire Naturelle, CNRS, Sorbonne Université, EPHE, Université Des Antilles, Paris, France

## Abstract

Although bacterial symbiosis is well-documented in chemosynthetic-based ecosystems, associations with microeukaryotes remain overlooked. In this study, using scanning electron microscopy and 18S rDNA barcoding, we investigate the presence, diversity, and biogeographic patterns of ciliate epibionts associated with two deep-sea caridean shrimp families: Alvinocarididae, and Thoridae. We identified a widespread lineage of ciliates colonizing the gills of different alvinocaridid species, extending their previously known distribution in freshwater and coastal habitats to deep-sea areas down to 3388 m. These ciliates form a distinct clade related to coastal Chonotrichia, but showing clear genetic divergence from the previously-described species. Geographic divergence of these ciliate populations was observed across the Pacific Ocean, with no evident structure related to their host species. These chonotrichian ciliates exhibited variation occurrence across host species, individuals, and regions, indicating a facultative association with their hosts. In contrast, the thorid shrimps harbored rare and phylogenetically diverse ciliates. More rarely, we found ciliates related to known parasitic lineages hosted by both shrimp families, with signs of immune response – black gills – in some individuals colonized by these ciliates. Our results reveal previously overlooked protist-crustacean associations in chemosynthetic ecosystems and highlight the ecological and biogeographic importance of this group in the deep ocean.

## Introduction

Deep-sea hydrothermal vent and hydrocarbon seep ecosystems are characterized by dense biological communities, dominated by animal species that are often specific to these habitats and found nowhere else (Van Dover 2000). These ecosystems are sustained by microbial communities through chemosynthetic primary production, unlike most others that rely on photosynthesis. A primary focus of hydrothermal vent and hydrocarbon seep research has been on both the visually dominant metazoans (e.g., aggregations of siboglinid tubeworms, bathymodioline mussels, and Rimicaris shrimps) and their chemosynthetic microbial symbionts (Dubilier et al. 2008; Sogin et al. 2021). In contrast, unicellular eukaryotes (protists) and parasitic associations have received much less attention in these environments. Despite this, they are recognized as key contributors to ecosystem functioning, serving as bacterial grazers, nutrient cyclers, and, in some cases, parasitic regulators of host populations (Hu et al. 2021). Free-living protists in hydrothermal vents are mostly dominated by the major clades of Alveolata and Radiolaria, as reported in most culture-independent studies (Hu et al. 2023; Murdock et Juniper 2019; Sauvadet et al. 2010). Early surveys of hydrothermal vent protists also revealed a high diversity of potential parasites within sediments, which are probably also hosted by the dense animal populations thriving around (López-García et al. 2003; Moreira 2003; Takishita et al. 2005). Conversely, in hydrocarbon seeps, protistan assemblages are mostly dominated by Radiolaria, Dinophyceae, and Cercozoa, while proportions of ciliates are comparatively lower (Li et al. 2024). These patterns indicate that microhabitats and fluid chemistry strongly influence the community composition and diversity of protists. Recent studies have also begun to investigate their functional role in chemosynthetic ecosystems and suggest that grazing activities from protists consume up to 28-62% of daily bacterial and archaeal biomass production in vent fluids (Hu et al. 2021). Similarly, in hydrocarbon seeps, protist predation and parasitism appear central to food webs and nutrient cycling (Li et al. 2024).

However, unlike macrofaunal communities, it remains contentious whether protist communities inhabiting vents and seeps consist mostly of cosmopolitan species or specialized taxa unique to these ecosystems (Murdock et Juniper 2019). Several studies have uncovered a wide diversity of novel protist lineages and taxa specific to a given vent locality (Coyne et al. 2013; Hu et al. 2023; Sauvadet et al. 2010) whereas others show that nearly 95% of protists from hydrothermal vents are also found in deep-sea background communities outside vent or seep influence (Murdock et Juniper 2019). These contrasting observations may be due to differences in sampling strategies and sample types (e.g., water, fluids, sediments, animal tissue), emphasizing the importance of microhabitats in determining their distribution. While clear evidence for vent-endemic protists remains rare, one notable exception are the colonial folliculinid ciliates forming dense “blue mats” on sulfide structures at hydrothermal vents (Kouris et al. 2007).

Symbiotic protist communities have been observed in host-associated niches in chemosynthetic environments, dominated by ciliates (Ciliophora), followed by Cercozoa and other lineages. These communities are found in the pallial cavity of the bathymodioline mussel *Bathymodiolus thermophilus* and the giant vesicomyid clam *Turneroconcha magnifica* from the vent fields in the Eastern Pacific (Sauvadet et al. 2010). These parasites may lead to mortality events observed in vent fauna (Moreira 2003). More broadly, this highlights the role of hydrothermal vent habitats as “oases” for protists. However, documented cases of protist-metazoan symbioses in chemosynthetic systems remain rare, with only one known example of an ectosymbiotic Syndiniales associated with mussels, *B. japonicus* and *B. platifrons* and the vesicomyid clams, *Calyptogena okutanii* and *C. soyoae*, at the Off Hatsushima seep in Sagami Bay, Japan (Noguchi et al. 2013).

This contrasts with other marine environments where numerous protist symbioses are well-documented and play a central role in structuring the entire ecosystems, such as *Symbiodinium* associations in coral reefs (LaJeunesse et al. 2018). Across marine environments, crustaceans also host a wide diversity of microeukaryotic symbionts, particularly ciliates, which belong mainly to three major ecological groups, apostome ciliates, chonotrichs, and suctorians, (Fernandez-Leborans et al. 2016). These groups differs in their life cycles, attachment strategies, and degrees of host specificity (Fernandez-Leborans et al. 2016). Among them, ectosymbiotic ciliates are observed on many groups including amphipods, isopods, decapods, and barnacles (Fernandez-Leborans et al. 2016). For instance, apostome ciliates commonly colonize gills of coastal crustaceans, generally without inducing strong host immune responses, although pathogenic associations such as black gill syndrome have reported in some penaeid shrimps (Fernandez-Leborans et al. 2013; Frischer et al. 2022; Landers et al. 2020). Chonotrichia ciliates are frequently found as gill epibionts of coastal and freshwater crustaceans, such as *Chilodochona carcini* on the mouthparts of the green crab *Carcinus maenas* (Fahrni et al. 2022; Lynn 2016).

Here, we explore the potential presence of similar ciliate symbionts associated with crustaceans from hydrothermal vents and hydrocarbon seeps, using two groups of deep-sea caridean shrimps as model cases. First is the family Alvinocarididae (Caridea: Bresilioidea) which are globally distributed and restricted to chemosynthetic ecosystems, with many species inhabiting vents and seeps in the Western Pacific (Methou, Chen, et al. 2024). The second is Thoridae (Caridea: Alpheoidea), specifically species of the genus *Lebbeus*. This taxon is broadly distributed across multiple oceanic regions, including the Pacific Ocean, North Atlantic, Southern Ocean, and Arctic regions, and isoften found in both chemosynthetic habitats as well as the ambient deep-sea environments (Komai et Chen 2024). Using a combination of scanning electron microscopy (SEM) and DNA barcoding (the nuclear 18S rDNA) to screen the gills of 11 alvinocaridid and 3 thorid shrimp species from chemosynthetic ecosystems across the Pacific, we address the following questions: 1) Are the gills of caridean shrimp from deep-sea chemosynthetic ecosystems colonized by epibiotic ciliates? 2) Are these ciliates also found in other marine environments, or are they preferentially associated with chemosynthetic habitats? 3) Is the phylogenetic diversity of these ciliates structured more by their geographic distribution or by the host shrimp lineages?

## Material and methods

### Sampling

We collected 14 shrimp species belonging to the Alvinocarididae (4 *Rimicaris* species and 7 *Alvinocaris* species) and Thoridae (3 *Lebbeus* species) families, from 21 chemosynthesis-based ecosystems across the Pacific Ocean, including 16 hydrothermal vent fields and five hydrocarbon seep sites at depths ranging from 628 m to 3388 m (Figure 1) (see Table S1 & S2 for additional details). Specimens were sampled using suction samplers mounted on various submersible vehicles, depending on the cruises, including the human-occupied vehicles (HOVs) including *Alvin, Nautile* and *Shinkai 6500*, or the remotely operated vehicles (ROVs) *Haima2, KM-ROV, ROPOS, SuBastian* and *Victor 6500*. Immediately after collection, specimens were preserved in 80% ethanol, fixed onboard directly in 2.5% glutaraldehyde for 16 hours, and stored in filtered seawater or directly stored at -80 °C. In the laboratory, individuals were examined morphologically under a stereomicroscope for species identification as well as for developmental stage (juvenile or adult) and sex (male, ovigerous, or non-ovigerous female). The body size of the shrimps was measured with a caliper to the nearest 0.1 mm. Gills were subsequently dissected and used for two types of analyses: microscopic observations and DNA extraction for molecular investigations of potential ciliate symbionts.

**Figure 1:**
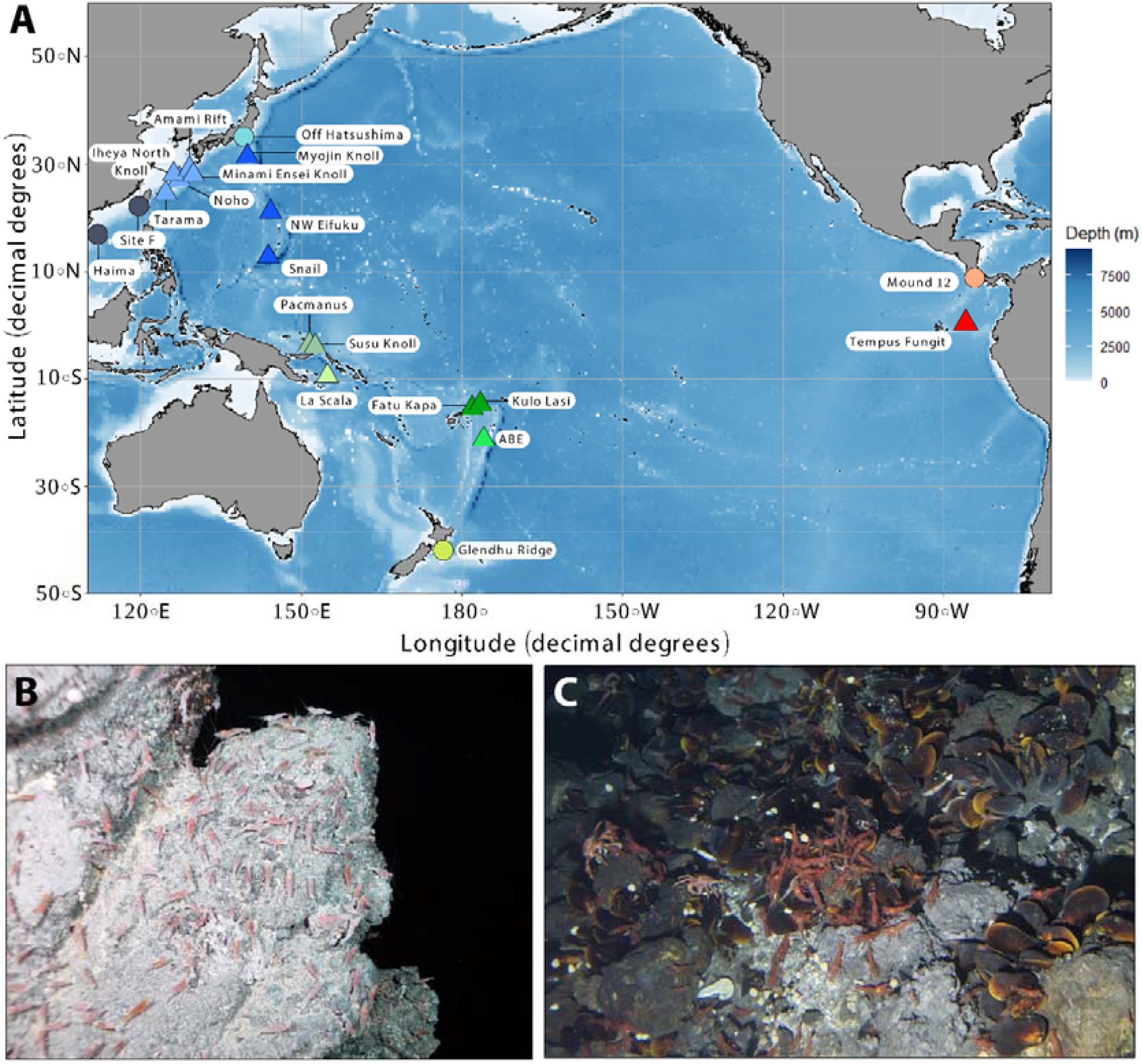
**A** Geographic map displaying hydrothermal vent (triangle) and hydrocarbon seep (circle) localities where shrimp specimens used in this study were collected. The map was generated in R (version 2024.04.2) with the ggOceanMaps package (v.2.2.0) (Vihtakari et al. 2024). Colors are consistent across figures and indicate the region where the samples were collected from: red (Eastern Pacific), green (Southwestern Pacific), and blue (Northwestern Pacific). **B** Alvinocaridid shrimps on sulfide rocks at the NW Eifuku vent field (Credit: Verena Tunnicliffe, University of Victoria, TN167 expedition) **C** *Lebbeus shinkaiae* shrimps gathering near mussel bed at the Minami-Ensei vent field (credit: YK23-16S expedition, JAMSTEC).

### Scanning electron microscopy (SEM)

For SEM observations, we primarily used specimens fixed onboard directly in 2.5% glutaraldehyde and rinsed in sterile seawater (Table S2). When on-board glutaraldehyde fixation was not available, we used specimens preserved in 80% ethanol or specimens that had been frozen at -80°C. In the latter case, the frozen specimens were post-fixed in 2.5% glutaraldehyde at the shore-based laboratory and subsequently rinsed three times with sterile seawater. In total, gills of 16 individuals were used for SEM observations, comprising 9 species of Alvinocarididae and 2 species of Thoridae. Dehydration was carried out for glutaraldehyde fixed specimens – either on board or at the laboratory – through an ascending ethanol series up to 100%. All samples, including those preserved in 80% ethanol, were then dehydrated up to 100% ethanol, followed by 5 h in a critical point dryer EM CPD 300 (Leica, Vienna) and coated with a thin layer of a 50/50 gold-palladium alloy using a Quorum Technologies SC7640 sputter coater (New Haven, USA). Observations were conducted using a Quanta 200 SEM (FEI-Thermo Fisher, Hillsboro, OR, USA). The microscope operated at an electron beam voltage of 5 to 10 kV, with magnifications ranging from 50× to 5,000×, depending on the structures examined.

### DNA Extraction, PCR, sequencing

DNA extractions were performed on 208 individual shrimps using one or more dissected gill filaments following the manufacturer’s instructions of the DNeasy PowerSoil Kit (Qiagen). Partial sequences of the 18S ribosomal gene were amplified by PCR using the ciliate-specific primers F384dT (5’-AAGGGCACCACCAGGAGTGG-3’) and R1147dT (5’-CGCTTTCGTGATYCTTGGTTCTC-3’) (Dopheide et al., 2008). PCR reactions were prepared in a 25 µL volume, comprising 15.25 µL of ultrapure water, 5 µL of Green GoTaq buffer, 2 µL of MgCl_2_, 1 µL of dNTPs, 0.75 µL of each primer, 0.25 µL of GoTaq Polymerase (Promega), and 1 µL of gill-derived DNA. Amplification conditions were as follows: initial denaturation at 94⍰°C for 5 min, followed by 40 cycles of 94⍰°C for 1 min, 55⍰°C for 1 min, and 72⍰°C for 2 min, with a final extension at 72⍰°C for 7 min. PCR efficiency was checked via 0.8% agarose gel electrophoresis. Samples with weak or faint amplification were subjected to nested PCR, using 1 µL of the first-round product as template under identical reagent and cycling conditions, except that the cycle number was reduced to 10 or 15 (vs. 40 cycles initially). Additionally, for some samples showing double bands on the agarose gel, gel extraction was performed on the band corresponding to the expected size, matching the one observed in samples with a single clear band. Extraction was carried out using the NucleoSpin® Gel and PCR Clean-up Kit (Macherey-Nagel).

Sequencing was performed using Sanger sequencing by Eurofins Genomics (Cologne, Germany). Raw chromatograms were cleaned, assembled, and edited using Geneious® 10.0.9 (https://www.geneious.com), removing artifacts, assembling fragments obtained with both primers for each individual. The resulting sequences were compared against the NCBI GenBank database using BLASTn (Basic Local Alignment Search Tool) for species identification, with the database version available as of December 2025.

### Phylogenetic reconstruction and haplotype network

Using BLAST results for a rough check of the taxonomic affinity, cleaned sequences were grouped by ciliate classes, resulting in two different alignments of sequences belonging to the Phyllopharyngea class (subclass Chonotrichia) and the class Oligohymenophorea (subclasses Apostomatia and Peritrichia). Alignment were conducted using the MUSCLE algorithm *(Edgar 2004)* implemented in Geneious and subsequently aligned together with reference ciliate sequences retrieved from the NCBI GenBank database, following (Lynn 2016) and (Fahrni et al. 2022) for the Phyllopharyngea alignment and (Metz et Hechinger 2021) for the Oligohymenophorea alignment. Phylogenetic reconstructions were performed using IQ-TREE v1.6.12 (Minh et al. 2020) with 1,000 ultrafast bootstrap replicates (Hoang et al. 2018). The best-fitting nucleotide substitution model for each tree was selected according to the Bayesian Information Criterion (BIC) using the ModelFinder module integrated in IQ-TREE. Resulting phylogenetic trees were visualized and edited using TreeViewer v2.0.2 (Bianchini et Sánchez-Baracaldo 2024). The 18S rDNA sequences of *Tokophrya huangmeiensis* (KJ567607) were used as outgroups to root the trees.

Given that the majority of ciliate 18S rDNA sequences obtained from alvinocaridid shrimp gills belong to a specific group within Chonotrichia, we conducted haplotype network analysis specifically on this group to examine their genetic structure. The alignment of the targeted sequences was performed using the previously mentioned methods, and the resulting alignments were utilized to construct a haplotype network using a median-joining method implemented in PopArt v1.7 (Leigh et Bryant 2015).

### Prevalence of ciliate epibionts

Shrimp individuals were grouped by subregion and by host species to consider both the influence of geography and host identity on the prevalence index. Given the recurrence of similar sequences belonging to Chonotrichia in our dataset, we calculated a prevalence index to estimate the frequency of gill colonization by Chonotrichae epibionts on our specimens. This index was calculated as follows:

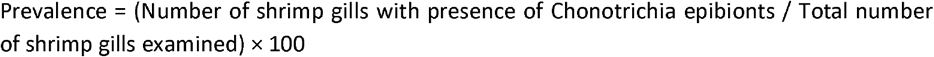

This yields the percentage of shrimp individuals colonized by Chonotrichia within each group. Individuals for which a 18S rDNA sequence affiliated to Chonotrichia was obtained by Sanger sequencing were considered as colonized by Chonotrichia epibionts. Conversely, the absence of colonization on individuals without 18S rDNA sequences or with sequences affiliated to another ciliate group was double confirmed by SEM or visual observation under a stereomicroscope using gills of the opposite side and floxin for enhancing the contrast between the ciliate epibionts and the shrimp tissue. For a few samples where microscopic observation was no longer possible, prevalence was assessed based solely on the barcoding results (Table S1).

## Results

### SEM observations of shrimp gill epibionts

Observations of gills by SEM confirmed the presence or absence of ciliates in several species of shrimps inhabiting chemosynthesis-based ecosystems. Colonization by these ciliates varied greatly in density among alvinocaridid shrimps, with some species exhibiting pronounced or even very dense colonization for most individuals observed in shrimp species such as *Alvinocaris longirostris* (Figure 2A & 2D), *Alvinocaris lusca* (Figure 2B & S2E), *Rimicaris vandoverae* (Figure 2C), and *Rimicaris variabilis* (Figure 2E & S2B). Other species like *Rimicaris cambonae*, however, showed limited colonization or no colonization at all (Figure 2F). There is also considerable variation within host species, as illustrated by the presence of both uncolonized and heavily colonized (Figure S2C and S2D) individuals of *A. longirostris* from the same locality (Off Hatsushima, Sagami Bay). Similarly, in *R. variabilis* specimens from one locality (Pacmanus, Manus Basin) were colonized (Figure S2B) while those from the other locality (La Scala, Woodlark Basin) was not (Figure S2A). In some individuals, such as *Rimicaris variabilis* (Figure 2E), filamentous bacteria were also observed on the gills and around the ciliates. No colonization by ciliates could be observed visually with our SEM observations on the gills of thorid shrimp species, neither *Lebbeus parvirostris* (Figure S1A) nor *Lebbeus shinkaiae* (Figure S1B).

**Figure 2:**
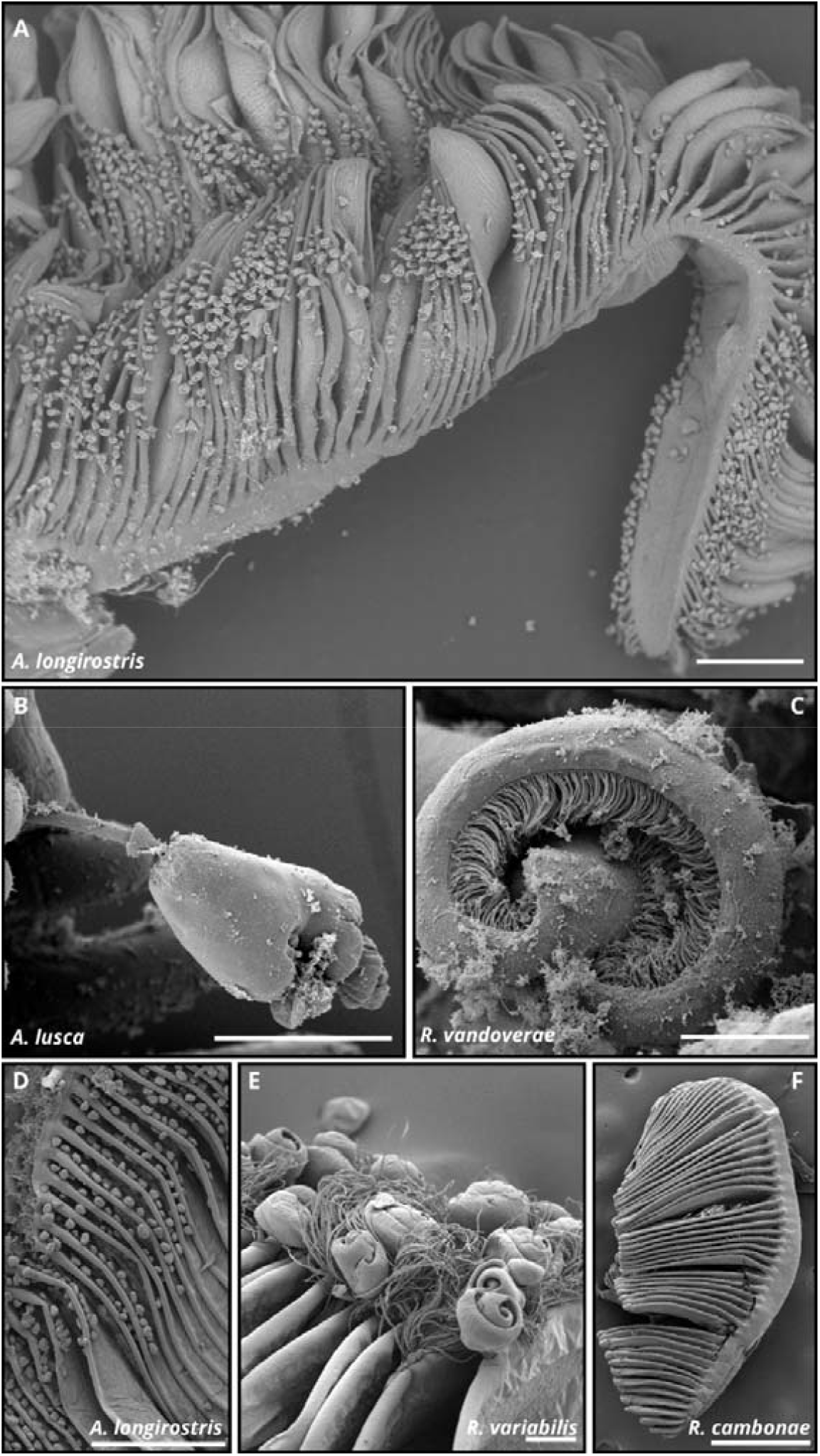
Scanning electron microscopy (SEM) observations of gills from multiple shrimp specimens collected from chemosynthesis-based ecosystems, either colonized or not by ciliates. **A :** Overview of an *Alvinocaris longirostris* gill filament colonized by several ciliates epibionts (500 µm), **C :** Close lateral view of a ciliate epibionts colonizing the gills of *Alvinocaris lusca* (50 µm), **B :** Close apical view of a ciliate epibionts colonizing gills of *Rimicaris vandoverae* (25 µm), **D :** View of a gill filament of *Alvinocaris longirostris* colonized by several ciliate epibionts (500 µm), **E :** View of several ciliate epibionts and the surrounding bacteria on a gill filament of *Rimicaris variabilis* (50 µm), **F :** Overview of a *Rimicaris cambonae* gill filament showing the absence of colonization by ciliate epibionts (500 µm).

The observed ciliates were approximately 50 µm in size, although variation could be observed, even within the same gill filament. They display a characteristic central invagination, surrounded by a well-defined ciliary arrangement forming a funnel-shaped collar composed of a coiled structure housing the cilia (Figure 2B). The body is elongate, and the ciliates are mainly localized at the base of the gill filaments, where they attach to the surface of the lamellae closest to the gill arch by means of a stalk (Figure 2C and 2A). This structure appears to serve as an anchoring support without penetrating the gill tissues (Figure S1C). Ciliates of various sizes can also be observed on the same gill (S1H).

### Phylogeny and 18S haplotype network of epibionts in Phyllopharyngea

Most 18S rDNA sequences of ciliates obtained from the gills of alvinocaridid shrimps (93 out of 120 sequences representing 18 unique sequences) were classified in Chonotrichia, a subclass within the class Phyllopharyngea (Figure 3). These sequences form a distinct monophyletic clade (bootstrap = 93) in our phylogenetic reconstruction, most closely related to *Chilodochona carcini* (GenBank Accession No.: KU588417) with a sequence similarity of 93.9 - 94.1%, or to *Spirochona gemmipara* (KY349104) with a sequence similarity of 94.3 - 94.6%. In contrast, a more limited number of 18S rDNA sequences from thorid *Lebbeus* shrimps were classified in Phyllopharyngea (4 out of 120 sequences representing three unique sequences). These sequences appear more scattered within our phylogenetic reconstruction of Phyllopharyngea, with one sequence (Phyllo_01 PX752129) isolated from *L. shinkaiae* most closely related to *Isochona* sp. (AY242116; 99.2 % similarity), and another sequence (Phyllo_03 PX752131) also isolated from *L. shinkaiae* being most closely related to *Acineta flava* (HM140400; 94.9% similarity). One sequence (Phyllo_02 PX752130) isolated from a gill of *L. unguiculatus* was affiliated with a group of sequences related to *Dysteria parasemilunaris voucher* (PP905720; 99.5% similarity). Only one sequence isolated from the gill of a *L. unguiculatus* individual (Chono_01 PX752096: see Figure 4B and Table S1) was affiliated with the group of sequences related to *Chilodochona carcini* (KU588417; 93.63% similarity).

**Figure 3:**
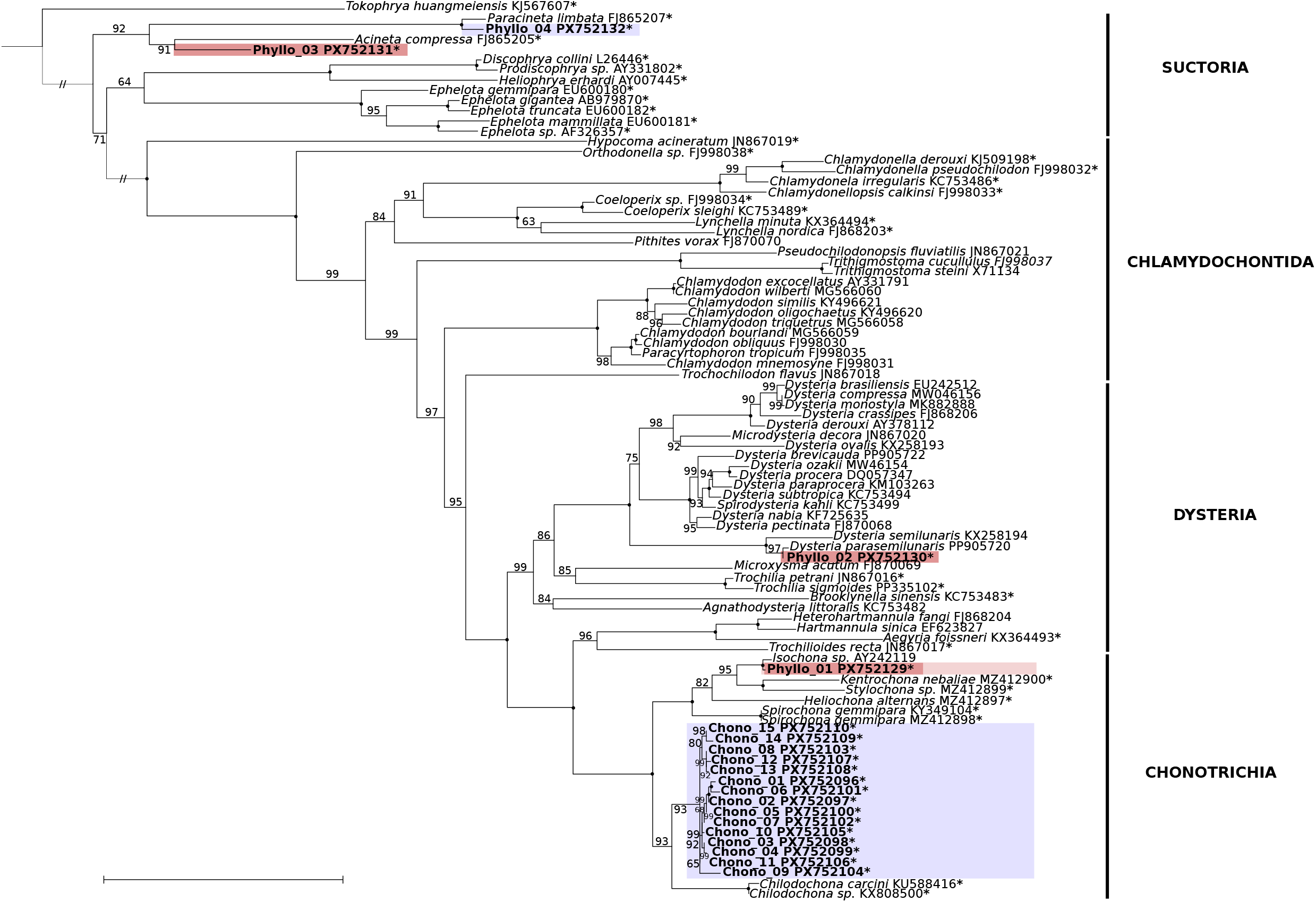
Phylogenetic tree of the Chonotrichae and related taxa (Phyllopharyngea class) on a partial fragment of the 18S ribosomal gene (length: 551 bp - 644 bp). The colors boxes indicate the family to which the ciliate host belongs, red: Thoridae, purple: Alvinocarididae. The asterisks indicate symbiotic ciliates associated with other metazoan hosts. The tree was inferred from SSU rDNA sequences using IQ-TREE v2.2.5 under the TIM2+F+I+R6 model selected by ModelFinder. NCBI accession numbers are shown to the right of each species name. The new sequences obtained in the present study are indicated in bold. Ultrafast bootstrap support values (10,000 replicates) are shown at the nodes, the ones with bootstrap of 100 are represented by black circles. The scale bar represents 0.05 substitutions per nucleotide position.

**Figure 4:**
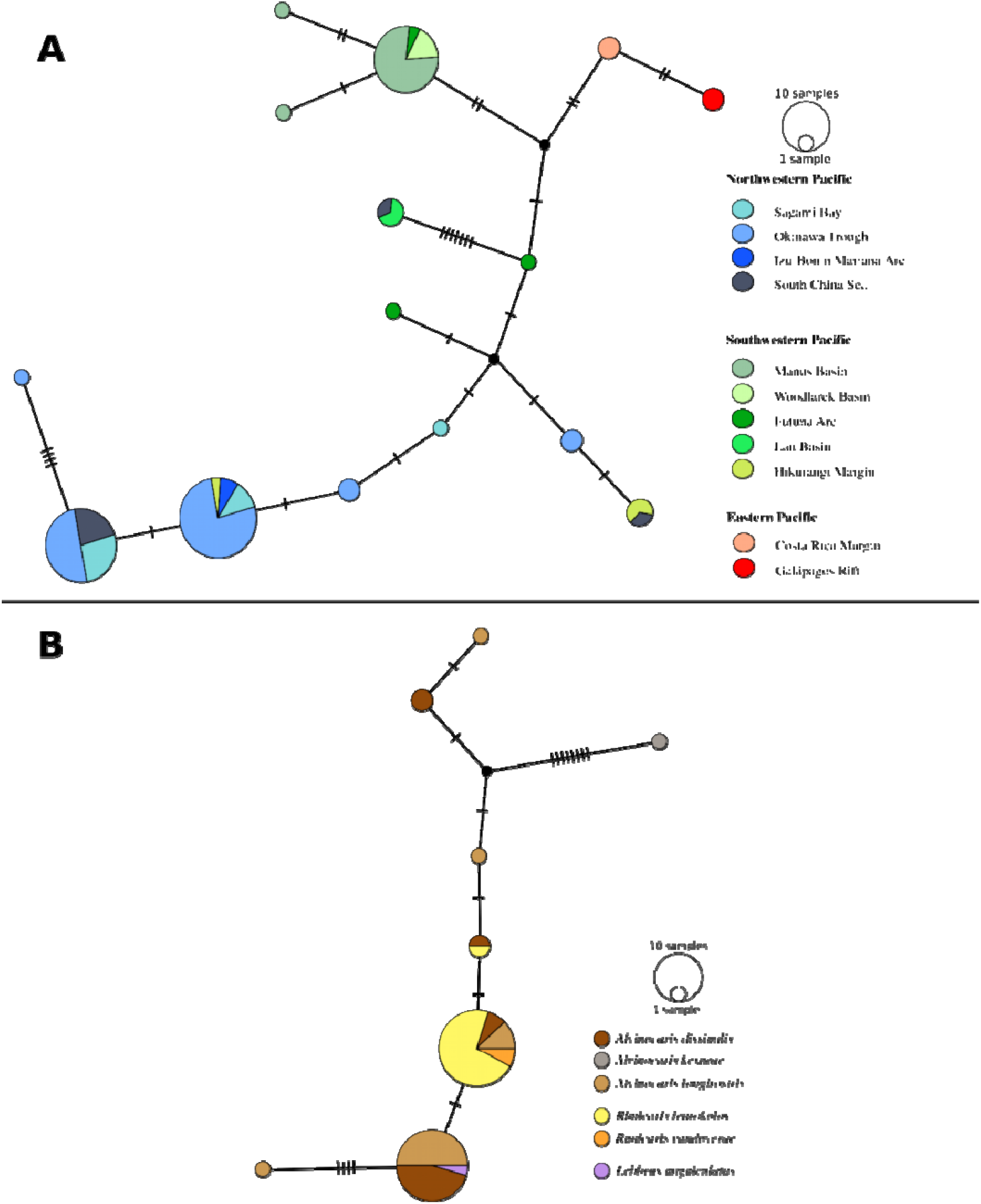
**A** Haplotype network based on a partial fragment of the 18S ribosomal gene (length: 551 bp - 644 bp)) of Chonotrichia ciliates according to geographic regions (86 individuals). Colors indicate the region of origin of the samples: red (Eastern Pacific), green (Southwestern Pacific), and blue (Northwestern Pacific). **B** Haplotype network based on the 18S ribosomal gene fragment of Chonotrichia ciliates according to phylogenetic host species of the North West Pacific (55 Individuals). Colors indicate the shrimp host species of the Chonotrichia ciliate. For both networks, each circle represents a distinct haplotype, with its size proportional to the number of sequences belonging to that haplotype. Lines connecting the circles represent mutations (nucleotide substitutions) between haplotypes. (see Table S3 & S4 for a detailed list of COI haplotypes and their corresponding GenBank ID).

Haplotype network of 18S sequences of *Chilodochona* spp. (Chonotrichia) shows that most haplotypes are structured by geographic localities (Figure 4A). In particular, Chonotrichia haplotypes from shrimp hosts inhabiting the Eastern Pacific were all unique haplotypes for their region, and most of the other haplotypes tended to cluster together between those from the Northwestern Pacific (Okinawa Trough, Sagami Bay, Izu-Bonin-Mariana Arc) and those from the Southwestern Pacific (Lau Basin, Futuna Arc, Manus Basin, Woodlark Basin). However, some Chonotrichia haplotypes contrasted with this general trend, like those isolated from shrimp hosts inhabiting the Hikurangi Margin in the Southwestern Pacific, which clustered with haplotypes from the Northwestern Pacific. Similarly, haplotypes from the South China Sea showed an intermediate position, with some sequences grouping with Northwestern Pacific clusters and others with Southwestern Pacific clusters. Finally, although we examined host identity as a potential structuring factor, we found no clear haplotype partitioning by host species (Figure 4B), indicating that host species does not appear to drive the geographic structuring observed here. This analysis was restricted to the Northwestern Pacific, as it was the only region in our dataset where multiple shrimp species coexist at the same location. Chonotrichia ciliates were detected in every subregion across all 11 study regions, indicating their ubiquitous distribution despite variations in abundance (Figure 1 & Table 1). All *Lebbeus* specimens (e.g., 3 species with a total sample size of 17) were removed from the prevalence analysis, as they were not colonized by Chonotrichia ciliates excepted one individual, which would have biased the colonization patterns observed. High prevalence values (≥50%), were found in 5 of the 11 regions studied, notably in Manus Basin vents (85%; n=20), Woodlark Basin vents (75%; n=4), the Sagami Bay seep (71%; n=14), Galápagos Rift vents (67%; n=3), and Costa Rica Margin seeps (50 %; n=6). Moderate prevalence values were found in the Okinawa Trough vents (46%; n=78) and South China Sea seeps (41%; n=17) whereas the Futuna Arc vents (27%, n=11), the Hikurangi Margin vents (25%; n=12), the Lau Basin vents (20%; n=10), and most notably the Izu-Bonin-Mariana Arc vents (10.5%; n=19) displayed low prevalence values. All shrimp species examined were colonized by Chonotrichia ciliates, except *Rimicaris cambonae*, for which no epibionts were observed.

**Table 1:**
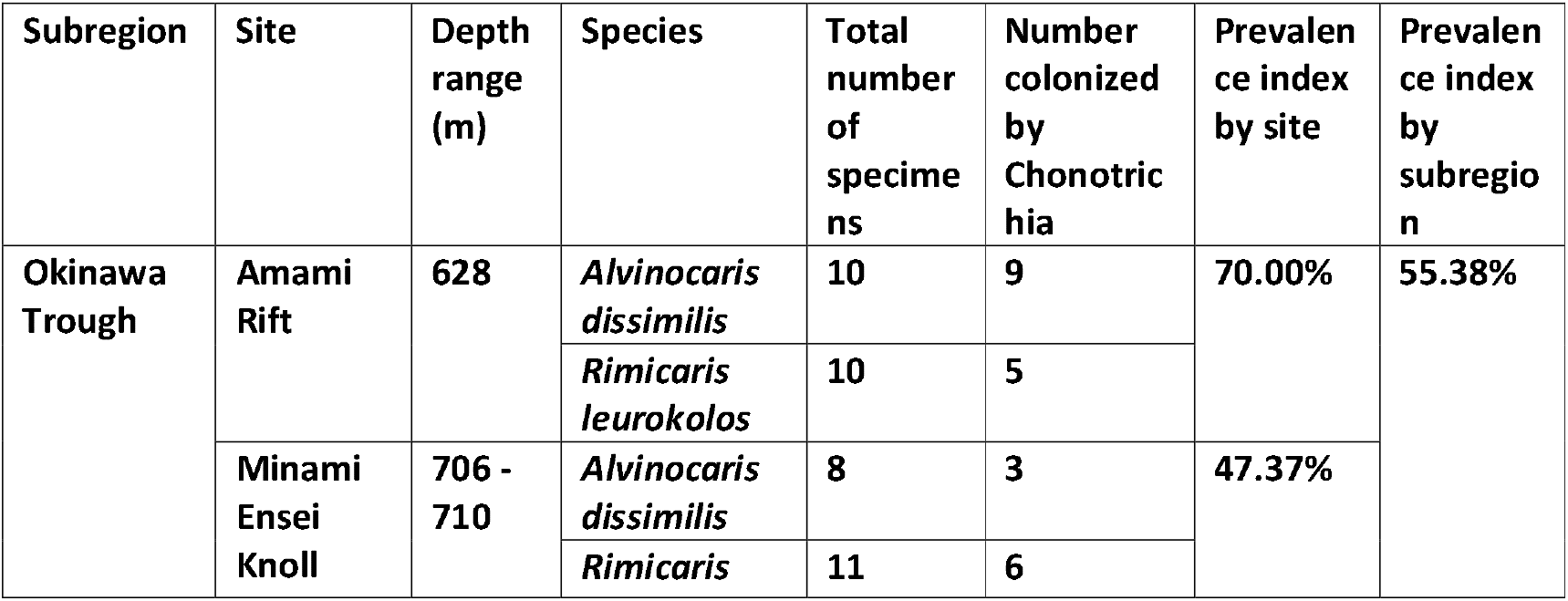
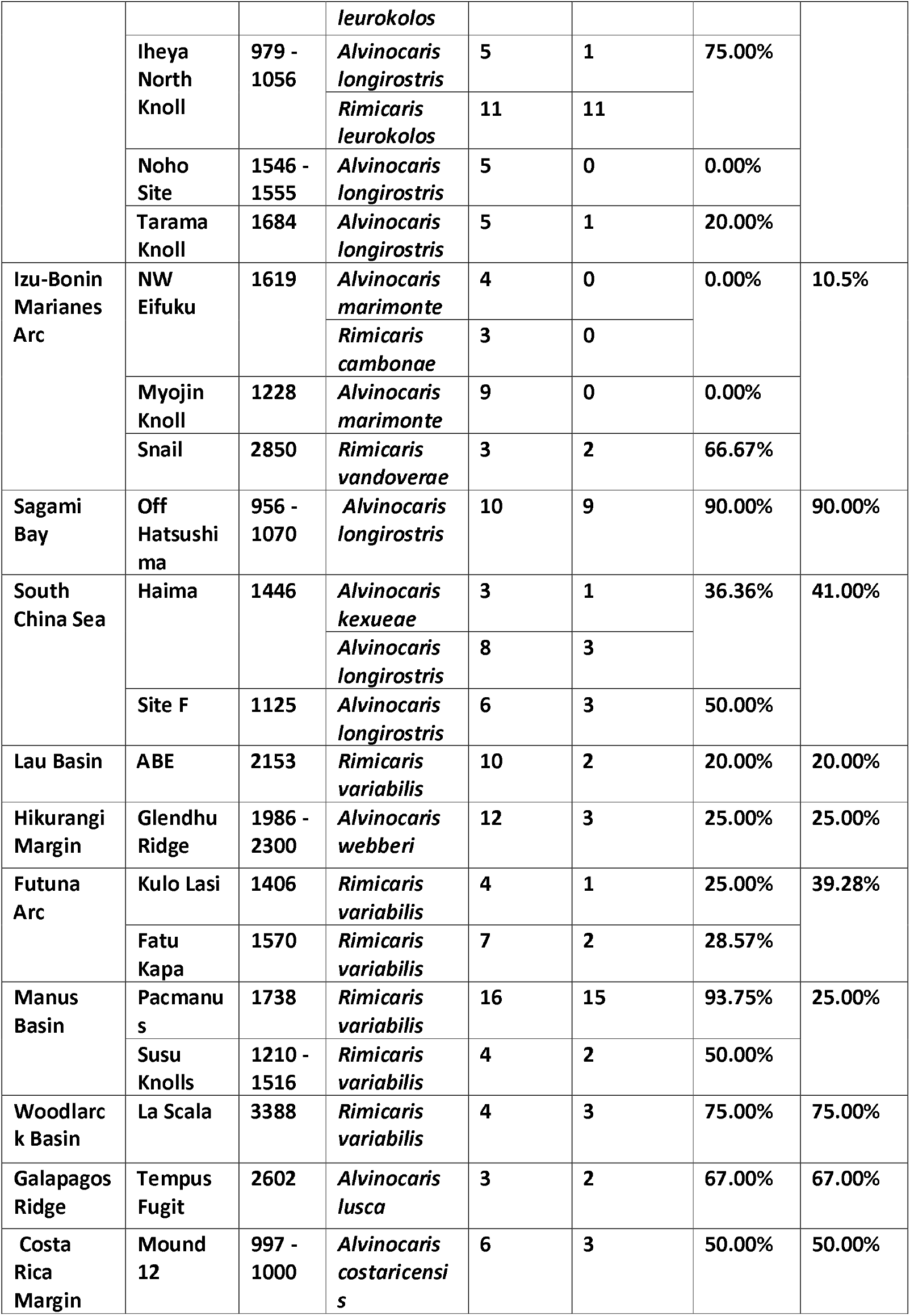
Prevalence of alvinocaridid individuals colonized by Chonotrichae ciliate epibionts.

### Phylogeny of ciliates symbionts in Oligohymenophorea

Although most 18S rDNA sequences from shrimp gills were classified as ciliates from the class Phyllopharyngea, some 18S rDNA sequences (23 out of 120 sequences representing 18 unique sequences) were affiliated with the class Oligohymenophorea, within the Apostomatida and Peritrichia groups (Figure 5). Most Apostomatida sequences were unique to a single shrimp host species, and identical ones came from individuals collected within the same region and sometimes from the same site. Hence, three sequences were shared among *R. variabilis* individuals from the Futuna Arc (Rva20, Rva29 & Rva30), two between *A. marimonte* from the Izu-Ogasawara Arc (Amt25 & Amt29), and three among *Lebbeus* spp. from the Okinawa Trough and Sagami Bay (Lpa1, Lpa2 & Lun14). Nearly all sequences from the subclass Apostomatia were positioned within three well-supported clades (bootstrap ≥ 90): one forming a sister branch to the group formed by the genera *Hyalophysa, Synophrya*, and *Vampyrophrya* (Oligoh_04 PX752114, Oligoh_03 PX752113 & Oligoh_05 PX752115), one basal node near genus *Metacollinia* (Oligoh_07 PX752117 & Oligoh_08 PX752118), and one basal branch sister to all Apostomatida comprising the rest of the sequences. One additional Apostomatida sequence (Oligoh_06 PX752116) showed the highest similarity to *Lynnia grapsolytica* (MT906161; 96.58% similarity). Three additional sequences were positioned within the subclass Colliniidae, with two sequences most closely related to *Pseudovorticella spathulata* (MH513617; 97.16% and 97.79% similarity) and one sequence (Oligoh_16 PX752126) most closely to *Rhabdostyla commensalis* (MN543651; 97.95% similarity).

**Figure 5:**
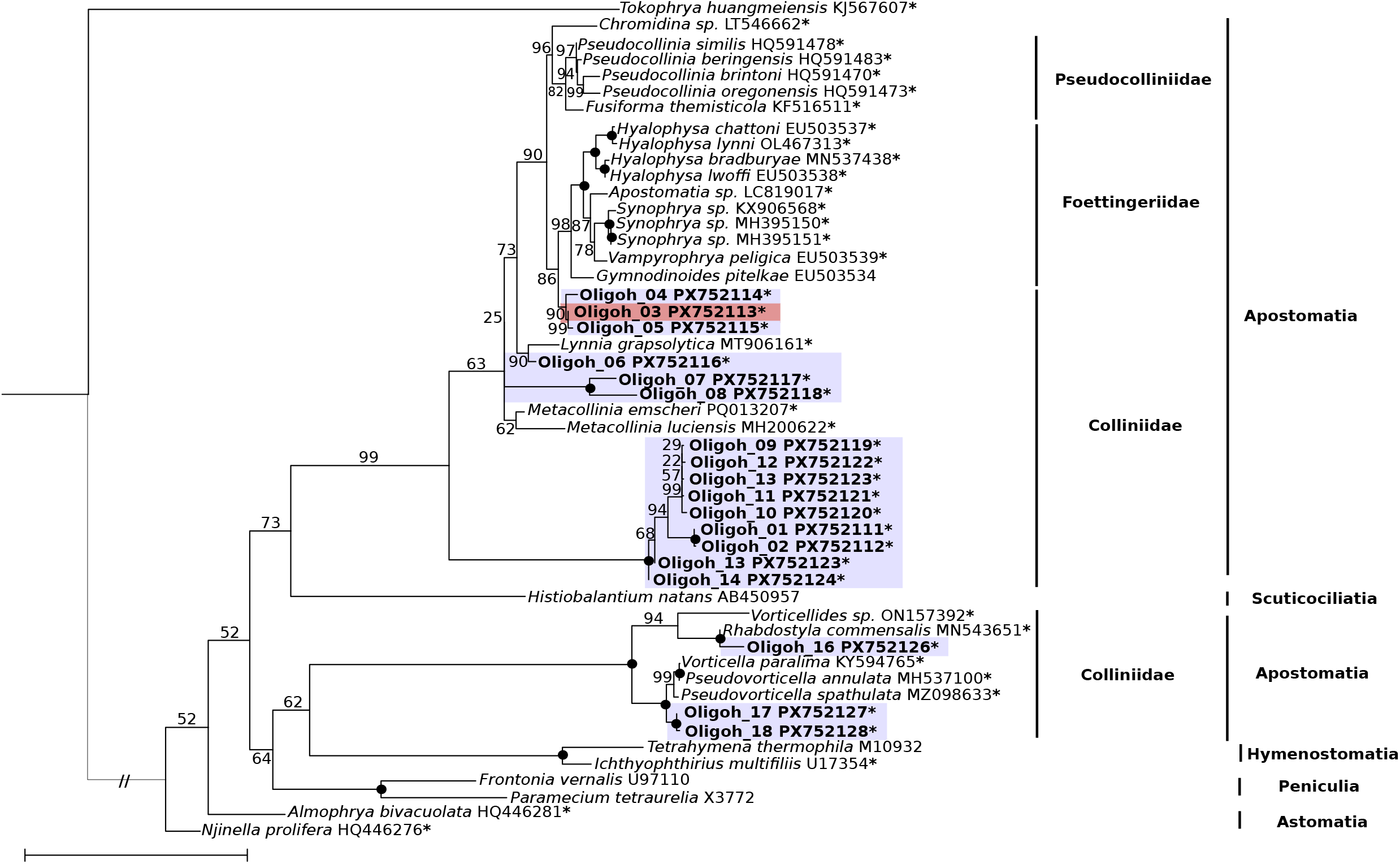
Phylogenetic tree of ciliates from the class Oligohymenophorea based on a partial fragment of the 18S ribosomal gene (length: 551 bp - 644 bp). The colors boxes indicate the family to which the ciliate host belongs, red: Thoridae, purple: Alvinocarididae. The asterisks indicate symbiotic ciliates hosted by other metazoan hosts. The tree was inferred from SSU rDNA sequences using IQ-TREE v2.2.5 under the TIM2+F+I+R3 model selected by ModelFinder. NCBI accession numbers are shown to the right of each species name. The new sequences obtained in the present study are indicated in bold. Ultrafast bootstrap support values (10,000 replicates) are shown at the nodes, the ones with bootstrap of 100 are represented by black circles. The scale bar represents 0.05 substitutions per nucleotide position.

Although systematic observations were not carried out for all these shrimp specimens, blackened gills were observed for some of the individuals from which an 18S rDNA sequence affiliated to Apostomatia was obtained, such as the *Rimicaris variabilis* specimen Rva20 (Oligoh_01; PX752111) from the Futuna Arc (Figure 6).

**Figure 6:**
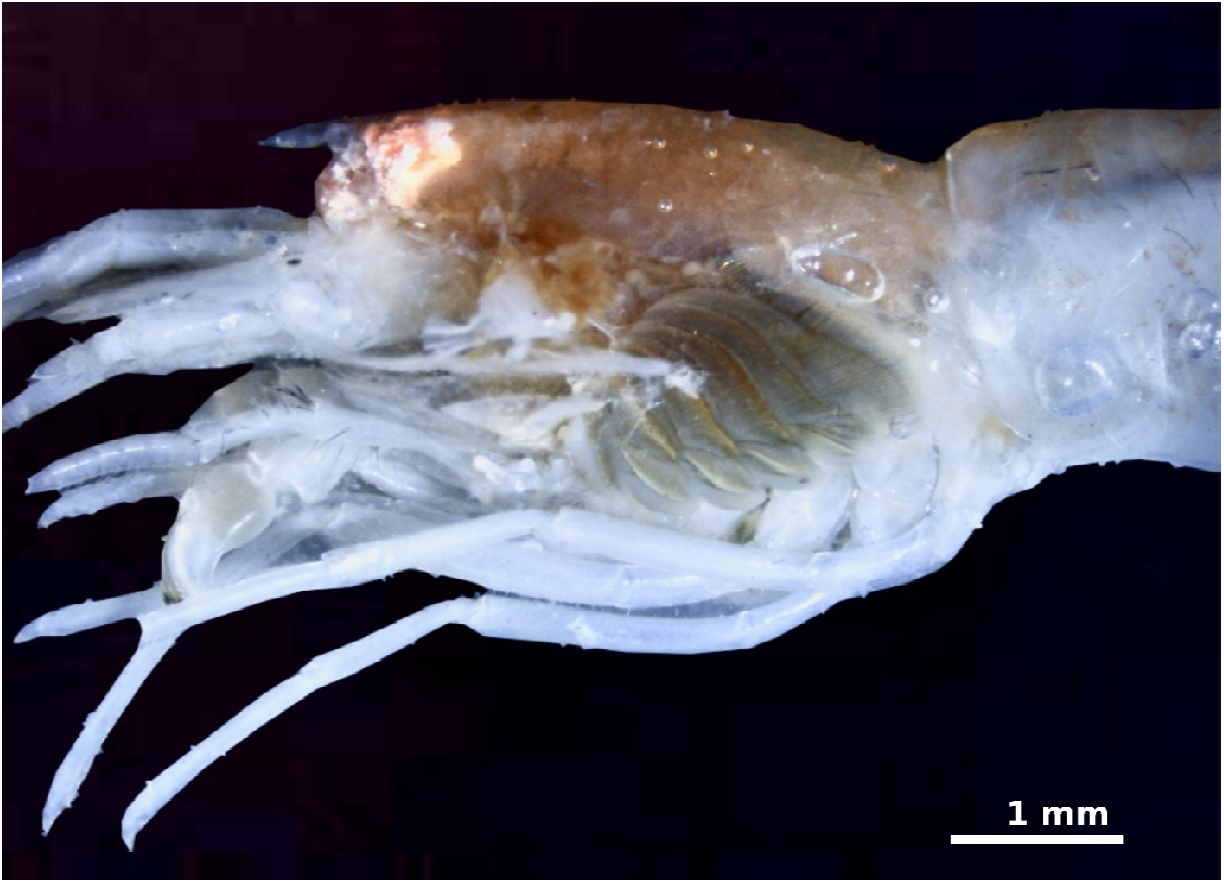
Photograph of a gill exhibiting black gill syndrome from an individual of *Rimicaris variabilis* (Rva20)

## Discussion

### A widespread lineage of Chonotrichae ciliate epibionts on gills of alvinocaridid shrimps

Epibionts of crustaceans belonging to Chonotrichia were previously reported in coastal or freshwater habitats (Lynn 2016; Fahrni et al. 2022). Our combined results from SEM (Figure 2) and molecular barcoding (Figures 3, 4 and 5) demonstrate that Chonotrichia ciliate epibionts are widely distributed on the gills of alvinocaridid shrimps from hydrothermal vents and hydrocarbon seeps throughout the Pacific Ocean (Figure 1), thus occupying deep and remote habitats down to 3388 m depth.

Our phylogenetic reconstruction places these ciliates as a sister group to *Chilodochona carcini* (Figure 3), a ciliate also known to live as an epibionts colonizing the green crab *Carcinus maenas* in coastal environments (Lynn 2016). Yet their relatively low similarity (93.9-94.1⍰#x0025;) and distinct morphological features to the closest known Chonotrichia implies the specificity of our newly detected lineage to deep-sea environments. The distinctiveness of this previously unknown Chonotrichia lineage must be balanced against the current under-representation of available reference sequences for this group, as highlighted by (Lynn 2016; Fahrni et al. 2022), with about 112 reported species but only 5 with associated molecular data. However, the specificity of Chonotrichia epibionts colonizing alvinocaridid shrimps is also supported by their host fidelity. Our examination of 208 host shrimp individuals shows that these ciliates were found almost exclusively on alvinocaridids, and not on the thorid shrimp genus *Lebbeus*, apart from one *L. unguiculatus* individual (Lun13), which was detected by 18S barcoding but could not be confirmed by SEM. Alvinocaridid shrimps are known only to inhabit chemosynthesis-based environments (Lunina et Vereshchaka 2014) – and even be restricted to hydrothermal vents for those belonging to the genus *Rimicaris* (Methou, Chen, et al. 2024)likely constraining their ciliate epibionts to the same habitat distribution. Unlike macrofaunal species, most studies on protist diversity have not been able to determine whether they are primarily specific to chemosynthetic environments or also cosmopolitan in deep-sea surroundings (Coyne et al. 2013; Hu et al. 2023; Murdock et Juniper 2019; Sauvadet et al. 2010). Even today, the colonial folliculinid ciliates forming dense “blue mats” in the Juan de Fuca Ridge are one of the rare cases of a vent endemic protist (Kouris et al. 2007). Our study reports another likely case of ciliates specific to chemosynthetic habitats: the Chonotrichia epibionts of alvinocaridid shrimps.

Given the observed variation in occurrence across host species, individuals, and regions, Chonotrichia epibionts appear to have a facultative association rather than an obligate relationship with their alvinocaridid hosts. Although the precise biological functions of these epibionts remain to be fully elucidated, their distribution patterns imply that they may play ecological roles comparable to those of closely related epibiotic ciliates. The size variations of ciliates observed on the same gill suggest continuous or progressive colonization, a process that may influence microbial turnover and interactions at the host surface (Bickel et al. 2012). It is likely that host characteristics such as cuticle structure, mucus composition, and immune responses play a crucial role in colonization success, in agreement with patterns observed in other epibiotic ciliates (Fokin 2004; Lynn 2010; Nowack et Melkonian 2010).

Unlike Alvinocarididae, colonization of thorid shrimp in the genus *Lebbeus* by ciliates differs considerably, with only a few individuals colonized each time by a very distinct lineage of Phyllopharyngea affiliated to *Acineta flava* (Suctoria), *Isochona* sp. sensu Snoeyenbos-West et al. 2004 (AY242116) (Chonotrichia), or *Dysteria paraselluminaris* (PP9005721). Only one symbiont affiliated to *Acineta* sp. sensu Gregori et al. 2016 (LK934647.1) was retrieved from one *Alvinocaris marimonte* shrimp, an alvinocaridid species that does not appear to be colonized by the specific Chonotrichae lineage. All these *Lebbeus* symbionts appear to belong to potentially new or existing lineages with 94.9% to 99.5% similarity to their closest known relative. However, unlike the epibionts specific to alvinocarids, it is difficult to determine whether these symbionts are specific to chemosynthetic environments, the deep ocean, or even cosmopolitan marine environments. Indeed, even though the host shrimp *Lebbeus shinkaiae* (Komai et al. 2012) and *L. parvirostris* (Komai et Chen 2024) have only been found in hydrothermal vents, other species within this genus, like *L. unguiculatus*, occupy other deep-sea habitats (Chan et al. 2010) and can be considered transient fauna dwelling in vents and seeps. Future studies on the associations of ciliates and other protists with a wider range of crustacean and invertebrate hosts will help to clarify these patterns of specificity to their hosts or their environments.

### Prevalence and biogeographic patterns of Chonotrichae epibionts

Although Chonotrichia epibionts frequently colonize gills of alvinocaridid shrimp species throughout the Pacific, we also observed considerable variability of their prevalence across sites and regions from 93% of the individuals colonized locally to a complete absence in a given population (Table 1). This prevalence reaches or exceeds 50%, for nearly half the sites, even though some values should be taken with caution due to a low number of specimens analyzed in some areas, particularly for the Eastern Pacific. In addition, some species, such as *Alvinocaris marimonte* (n = 14) and *Rimicaris cambonae* (n= 9), never appear to host Chonotrichia epibionts on their gills (Figure 2). Thus, although widespread, the presence of these ciliates varies among different species but also among individuals of the same species, indicative of a facultative association for their host.

The observed inter- and intra-specific variability in colonization of Chonotrichia epibionts among shrimp host species and localities is somewhat unclear even in the light of our results. Part of the inter-individual variability could be related to the molting cycle of alvinocaridids throughout their adult life, renewing their cuticle and probably the ciliate epibionts (Corbari 2008). While the complete absence of these epibionts in certain areas (e.g., NW Eifuku or Myojin Knoll; see Table 1) may be linked to local conditions of vent and seep geofluids, observations from the Okinawa Trough present a contrasting case. There, Chonotrichia epibionts colonize all examined alvinocaridid species across the geochemical spectrum, from hosts like *Rimicaris leurokolos* living near the hottest fluids to those like *Alvinocaris dissimilis* inhabiting cooler mussel beds (Yahagi et al. 2015). This pattern implies that colonization may depend more on host-specific factors than on local chemistry.

Our data also suggest a global geographic structuration of Chonotrichia, with a slight differentiation between large regional assemblages from the Northwestern, the Southwestern, and the Eastern Pacific (Figure 4A). This geographic structuration corresponds approximately to the biogeographic provinces defined for macrofaunal species at hydrothermal vents (Moalic et al. 2012; Tunnicliffe et al. 2024) and their connections with hydrocarbon seeps (He et al. 2023). Nevertheless, some exceptions to this geographic pattern are notable. For instance, ciliate epibionts from shrimp hosts off New Zealand show stronger genetic similarities with those from the Northwestern Pacific despite more than 7000 km of distance. This genetic similarity between distant regions mirrors the one observed for their alvinocaridid hosts, like *Alvinocaris dissimilis*, whose distribution range extends from the Okinawa Trough near Japan to the Hikurangi Margin in New Zealand (Methou et al. 2024). Additionally, methane seeps from the South China Sea appear to function as a corridor that links the Northwestern and the Southwestern Pacific regions (He et al. 2023). Indeed, distant populations of their alvinocaridid host *A. longirostris*, which are present in both the South China Sea and Sagami Bay seeps, the Okinawa Trough vents, and as far south as the Manus Basin vents, have also been shown to exhibit similar patterns of gene flow and population structure (Dai et al. 2025). This broad distribution, which stretches from Manus Basin to the Okinawa Trough, suggests that the South China Sea may encourage multi-step dispersal events across basins, which may also enable the associated epibionts to be transported over long distances. The sharing of similar hosts between regions could thus facilitate long-distance transport of their epibionts (Goffredi et Orphan 2010). Nevertheless, it should be noted that the usage of the relatively conserved 18S rRNA gene marker may be limited in its ability to resolve fine-scale geographic structure among the different Chonotrichia epibionts populations.

### Other infrequent and potentially parasitic ciliates belonging to Oligohymenophorea

In addition to the dominant Chonotrichia epibionts, ciliates belonging to Oligohymenophorea could also be found, although more sporadically, notably with some Peritrichia (*Vorticella* or *Pseudovorticella*) or some Apostomatia (*Metacollinia, Hyalophysa, Pseudocollinia, or Lynnia* among others) (Figure 5). Their presence is more geographically restricted than that of Chonotrichia epibionts, most often within a single region, and for some of them, it could correspond to a parasitic relationship. This is supported by the immune response observed in an individual of the alvinocaridid shrimp *R. variabilis* (Rva20) displaying blackened gills (Figure 6). This specimen is colonized by an apostome ciliate most closely related to *Hyalophysa*, a genus known to parasitize crustaceans in coastal environments (Frischer et al. 2022). Some species of this genus, such as *Hyalophysa lynni*, have been reported to induce physiological imbalances and trigger an immune response characteristic of black gill syndrome (Bickel et al. 2012; Fernandez-Leborans et al. 2013; Frischer et al. 2022; Landers et al. 2020). Although no direct tissue damage was observed in the Rva20 individual, the presence of *Hyalophysa* and the associated blackened gill coloration typical of an immune response suggests a potential physiological impact that has not been observed in individuals colonized by Chonotrichia, either in coastal shrimps (Frischer et al. 2022) or in deep-sea alvinocaridids examined in this study.

It is plausible that some hydrothermal vent areas, like the Fatu Kapa site on the Futuna Arc, or some periods, could be more affected than others by these potential parasites. In coastal penaeids, severe outbreaks of black gill disease have particularly affected areas off the coasts of South Carolina and Georgia, compared to other regions (Frischer et al. 2022; Landers et al. 2020). Seasonality also appears to play an important role in these epidemics, with the presence of black gills and *Hyalophysa lynni* parasites mostly in late summer when water temperatures reach their maximum and oxygen levels are at their lowest (Frischer et al. 2022). Although seasonal dynamics and biological cycles are structured differently in the absence of light in deep-sea chemosynthetic ecosystems (Mat et al. 2020; Methou et al. 2022; Zhang et al. 2025), black gill syndrome in alvinocaridid shrimps could also follow periodic bursts throughout the year.

Our results align with the conclusions of (Dykman et al. 2023), who proposed that the ecological isolation and unstable nature of hydrothermal vent environments do not significantly limit the diversity of metazoan parasites, particularly trematodes, but also copepods or nematodes. Contrary to previous expectations, the high density of animals within these ecosystems could instead promote the transmission and maintenance of parasites and epibionts (Moreira 2003).This diversity also includes parasites with indirect life cycles that involve multiple hosts (L. Dykman et al. 2025). Our work expands this diversity of potential parasites to microeukaryotes with the example of alvinocaridid shrimps displaying immune blackened gill responses when infected by Apostomatia ciliates related to *Hyalophysa*. Our study has only scratched the surface of the many potential relationships between ciliates and animals living side-by-side in deep-sea chemosynthetic ecosystems, and we believe that many more remain to be discovered.

## Acknowledgements

We are indebted to the captain and crew of R/Vs *Atlantis* (cruise AT42-03) *Atalante* (cruises Futuna1: https://doi.org/10.17600/10010110; Futuna3: https://doi.org/10.17600/12010040; and Chubacarc: https://doi.org/10.17600/18001111) *Yokosuka* (cruises YK10-11, YK17-17, YK23-06, and YK23-16S), *Kairei* (cruise KR15-17), *Kaimei* (cruise KM23-E05), *Tangaroa* (cruise TAN2102), and *Falkor (too)* (cruise FKt231024), *Tan Kah Kee* (Site F cruise), and *Haiyang Dizhi* 6 (Haima cruise) for their excellent support for our scientific activities; we extend the same gratitude to pilots and technical teams of the HOV *Alvin, Nautile, Shinkai 6500* and ROVs *Haima2, Kaiko* (vehicle *Mk-IV*), *KM-ROV, ROPOS*, and *SuBastian* and *Victor 6000*. Chief scientists of the relevant research cruises are also gratefully acknowledged for their efforts in organizing and executing the expeditions: Erik Cordes (Temple University) for AT42-03, Didier Jollivet (CNRS) and Stéphane Hourdez (CNRS) for Chubacarc; Yves Fouquet (Ifremer) for Futuna1 & Futuna3, Shigeaki Kojima (Atmosphere and Ocean Research Institute, the University of Tokyo) for YK10-11, Hiroko Makita (Tokyo University of Marine Science and Technology) for YK17-17, Masahiro Yamamoto (JAMSTEC) for YK23-06, Kenichiro Tani (National Museum of Nature and Science, Tsukuba, Japan) for YK23-16S, Shinsuke Kawagucci (JAMSTEC) for KR15-17, Ken Takai (JAMSTEC) for KM23-E05, Laura Wallace (New Zealand Institute of Geological and Nuclear Sciences) for TAN2102, John W. Jamieson (Memorial University of Newfoundland) for FKt231024, Dr. Huaiyang Zhou (Tongji University) for the Site F cruise, and Dr. Jun Tao (Guangzhou Marine Geological Survey (Guangzhou, China) for the Haima seep cruise. Sampling during cruise KM23-E05 within the Islands Unit of the Mariana Trench Marine National Monument of the United States of America was carried out under the special use permit SUP 12542-23001 (MSR U2022-047). We thank Hidetaka Nomaki (JAMSTEC) for ship time and seagoing support, and Shinsuke Kawagucci (JAMSTEC) for leading and drafting the proposal for the R/V *Kaimei* cruise KM23-E05 with Ken Takai. We thank Nicolas Gayet (UMR BEEP; Ifremer) for his work in preparing samples for scanning electron microscopy. We also extend our gratitude to Prof. Pei-Yuan Qian (The Hong Kong University of Science and Technology), Prof. Jian-Wen Qiu (Hong Kong Baptist University, HKBU), Dr. Yi-Tao Lin (HKBU), Ms. Qi Dai (HKBU), and Dr. Yi-Xuan Li (HKBU) for their contributions to collecting and processing samples from the South China Sea seeps.

## Funding

Lison Hey and Pierre Methou were supported by ISblue project, Interdisciplinary graduate school for the blue planet (ANR-17-EURE-0015) and co-funded by a grant from the French government under the program “Investissements d’Avenir” embedded in France 2030. Analyses conducted during this study also benefited from French State aid managed by the National Research Agency under France 2030 LIFEDEEPER: ANR-22-POCE-0007. Dewi Langlet was supported by JSPS KAKENHI grant 23K05942. This study was also supported by two General Research Funds (GRFs) and one Collaborative Research Fund (CRF) of the Hong Kong SAR government (16309324, 16100425, and C2013-22G), a project from the Otto Poon Center for Climate Resilience and Sustainability of The Hong Kong University of Science and Technology (CCRS25SC01), and two projects from the Southern Marine Science and Engineering Guangdong Laboratory (Guangzhou) (2021HJ01 and SMSEGL24SC01). The R/V *Yokosuka* research cruise (YK23-16S) was supported by the Cooperative Research Program of Atmosphere and Ocean Research Institute, The University of Tokyo (R/V *Yokosuka*, cruise YK23-16S). The R/V *Kairei* research cruise KR15-16 was supported by Council for Science, Technology, and Innovation (CSTI) as the Cross Ministerial Strategic Innovation Promotion Program (SIP), Next-generation Technology for Ocean Resource Exploration. The R/V *Falkor (too)* cruise FKt231024 (Project Zombie: Bringing dead vents to life – Ultra fine-scale seafloor mapping”) was funded by the Schmidt Ocean Institute. The AT42-03 expedition off Costa Rica was funded by the NSF grant NSF OCE 1635219.

## Notes

### Competing Interest Statement

The authors have declared no competing interest.

