## Supplementary Figure S1 to S3 for "Widespread symbiosis of ciliate epibionts colonizing gills of shrimps inhabiting vents and seeps across the Pacific Ocean"

### Supplementary Material

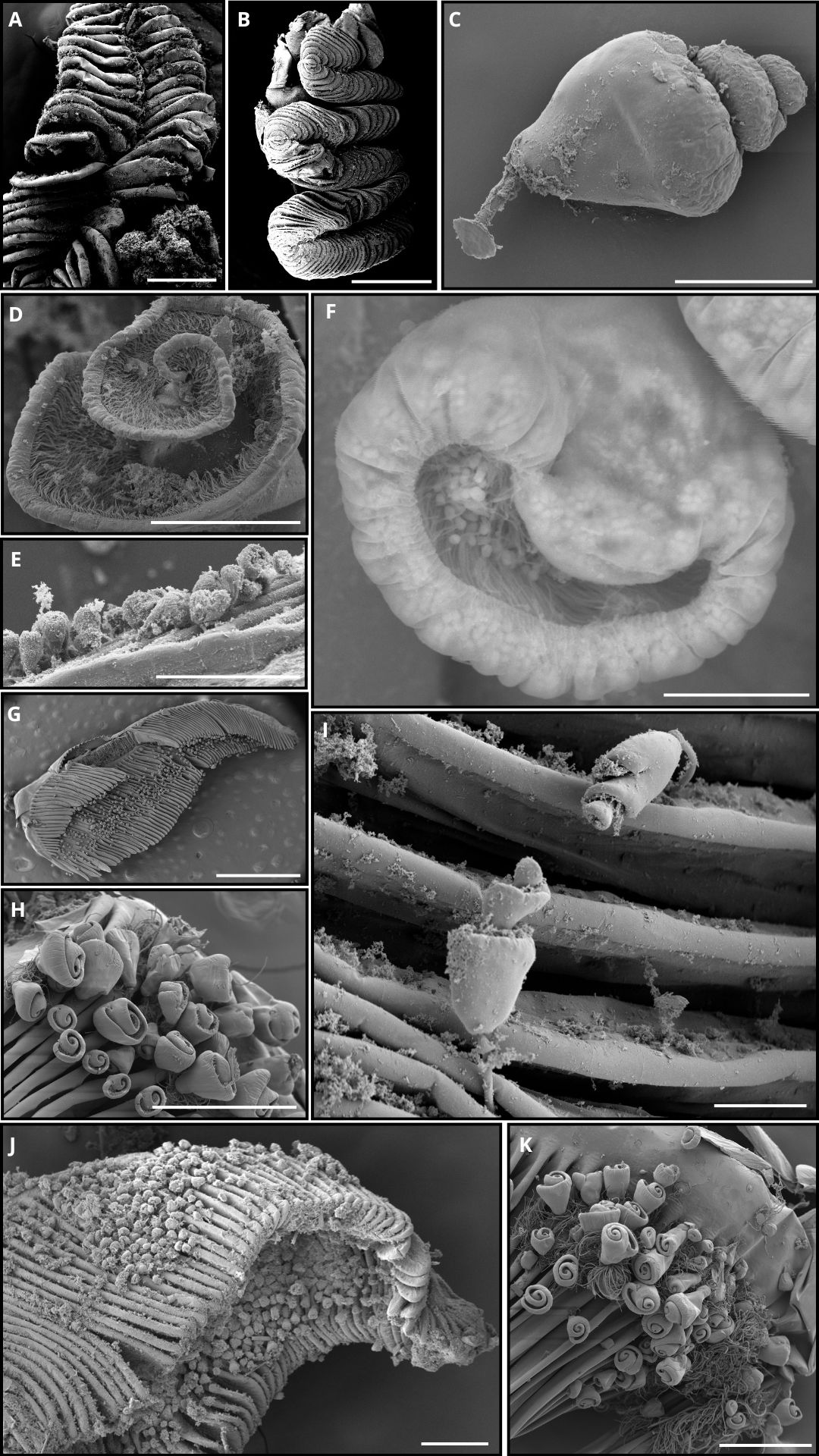

**Figure S1** : Scanning electron microscopy (SEM) observations of gills from multiple shrimp specimens collected from chemosynthesis-based ecosystems, either colonized or not by ciliates. (**A** : Overview of a gill filament of *Lebbeus parvirostris* showing no colonization by epibiotic ciliates (200 µm), **B** : Overview of a gill filament of *Lebbeus shinkaiae* showing no colonization by epibiotic ciliates (1.5 mm), **C** : Close-up of an epibiotic ciliate that colonized a gill filament of *Alvinocaris longirostris* (100 µm), **D** : Close-up of an epibiotic ciliate colonizing a gill filament of *Alvinocaris longirostris* (50 µm), **E** : Epibiotic ciliates colonizing the gills of *Rimicaris leurokolos* (200 µm), **F** : Close-up of an epibiotic ciliate colonizing the gills of *Alvinocaris longirostris*, showing bacteria inside (15 µm), **G** : Overview of a gill filament of *Alvinocaris longirostris* colonized by epibiotic ciliates (1.5 mm), **H** : Overview of a gill filament of *Rimicaris variabilis* colonized by epibiotic ciliates (200 µm), **I** : View of an epibiotic ciliate on the gill lamellae of *Alvinocaris costaricensis* (50 µm), **J :** Overview of an epibiotic ciliate colonizing the gills of *Alvinocaris dissimilis* (250 µm), **K :** View of epibiotic ciliates colonizing the gills of *Rimicaris variabilis* along with surrounding bacteria (200 µm))

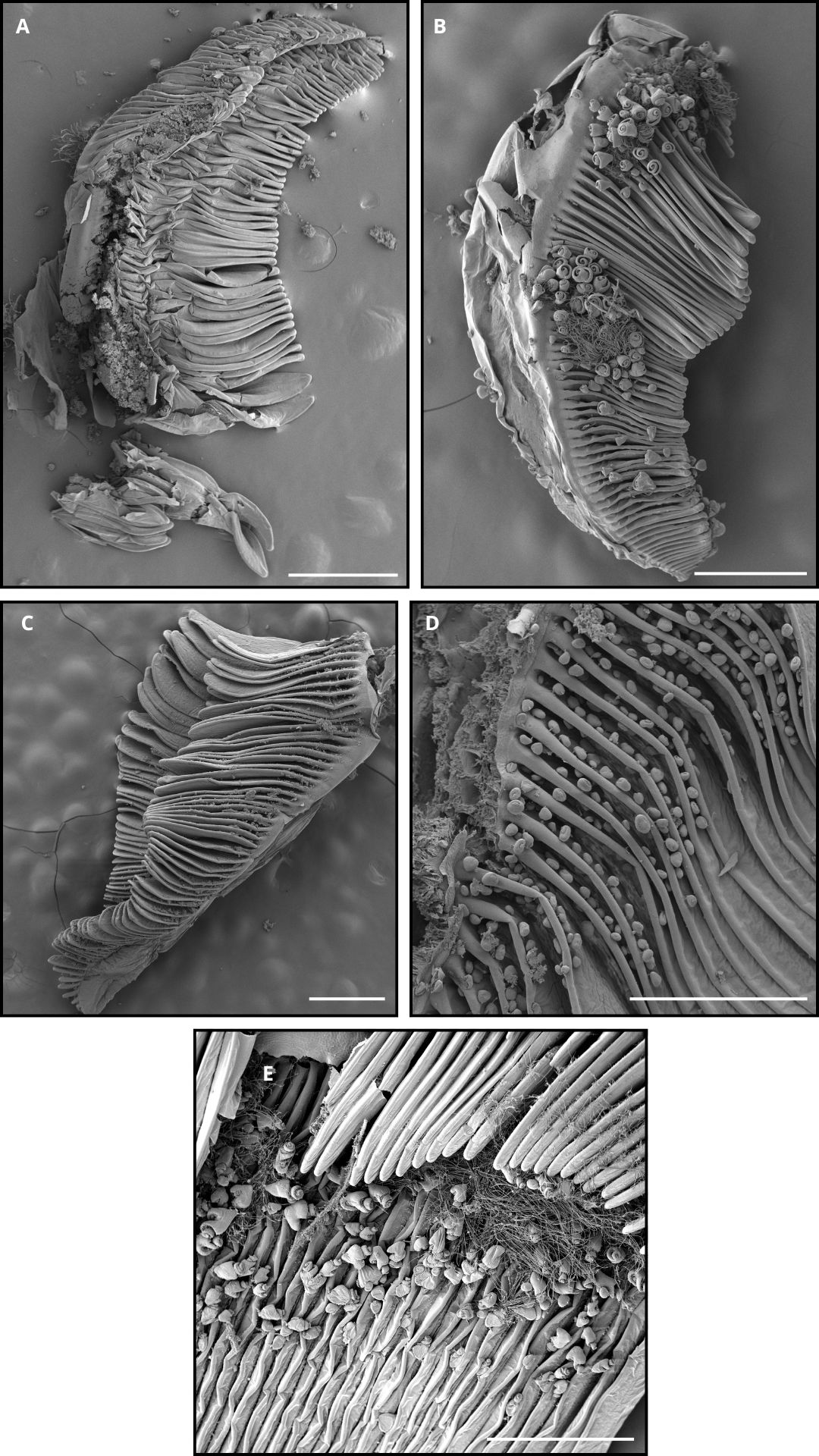

**Figure S2** : Scanning electron microscopy (SEM) observations of gills from multiple shrimp specimens collected from chemosynthesis-based ecosystems, either colonized or not by ciliates. **A** : Overview of a gill filament of *Rimicaris variabilis* showing absence of colonization by ciliates (500 µm), **B** : Overview of a gill filament of *Rimicaris variabilis* showing presence of colonization by ciliates and bacteria (500 µm), **C** : Overview of a gill filament of *Alvinocaris longirostris* showing absence of colonization by epibiotic ciliates) (500 µm), **D** : Overview of a gill filament of *Alvinocaris longirostris* showing presence of colonization by epibiotic ciliates (500 µm), **E** : Overview of a gill filament of *Alvinocaris lusca* colonized by epibiotic ciliates (500µm))
